# Enabling the democratization of the genomics revolution with a fully integrated web-based bioinformatics platform

**DOI:** 10.1101/040477

**Authors:** Po-E Li, Chien-Chi Lo, Joseph J. Anderson, Karen W. Davenport, Kimberly A. Bishop-Lilly, Yan Xu, Sanaa Ahmed, Shihai Feng, Vishwesh P. Mokashi, Patrick S. G. Chain

## Abstract

Continued advancements in sequencing technologies have fueled the development of new sequencing applications and promise to flood current databases with raw data. A number of factors prevent the seamless and easy use of these data, including the breadth of project goals, the wide array of tools that individually perform fractions of any given analysis, the large number of associated software/hardware dependencies, and the detailed expertise required to perform these analyses. To address these issues, we have developed an intuitive web-based environment with a wide assortment of integrated and cutting-edge bioinformatics tools. These preconfigured workflows provide even novice next-generation sequencing users with the ability to perform many complex analyses with only a few mouse clicks, and, within the context of the same environment, to visualize and further interrogate their results. This bioinformatics platform is an initial attempt at Empowering the Development of Genomics Expertise (EDGE) in a wide range of applications.

## INTRODUCTION

The field of genomics has made tremendous technological leaps in recent years, and the combined decrease in sequencing costs and expansion in applications (transcriptomics, metagenomics, single cell genomics) have truly revolutionized the way scientists approach biological questions (for a recent review, see (Buermans and den Dunnen 2014)). Now that a trained technician can single-handedly produce gigabases of sequence data in essentially a day’s work, “next generation sequencing” (NGS) is being applied by many smaller laboratories, as well as the large traditional sequencing centers, across a wide range of disciplines in order to answer a variety of complex problems. For instance, NGS is being applied to the characterization and attribution of outbreaks in clinical environments (Conlan et al. 2014), food safety (den Bakker et al. 2014), the development of alternative energy sources (Wang et al. 2012; Wohlbach et al. 2014), and many other fields.

Although many advances have been made in bioinformatics methods development, the so-called “democratization of genomics” (Koren et al. 2014) has not yet fully expanded to the bioinformatic realm, making it difficult for investigators to adequately analyze genomic big data (Daber et al. 2013; Watson-Haigh et al. 2013). While NGS no longer seems new, it has really only been since 2005 that a revolutionary new technology (pyrosequencing) (Margulies et al. 2005) was introduced after more than twenty years of chemical degradation (Maxam and Gilbert 1977) and chain termination (Sanger et al. 1977) sequencing. Some of these NGS technologies have already been abandoned even after strong market performance; other new technologies are only now emerging, and the ones that have thus far survived continue to undergo improvement. Despite reads of limited length, Illumina^®^ (Bennett 2004) currently dominates the market, in part due to its very high throughput and low cost.

Analysis of the massive datasets produced in NGS studies and interpretation of the results requires expertise in both computer science and biology. Therefore, although the decreasing cost and decreasing laboratory footprint of NGS technologies make the production of these datasets a more realistic goal for many laboratories, there still remain at least three core issues in bioinformatics that hamper the broader use of NGS data. First, the numerous and diverse specific questions being asked of NGS data require highly specialized pipelines. While any given question can sometimes make use of the same basic tool(s) with different parameters and post-processing, other questions may require similar bioinformatic manipulation but are optimally answered using different tools, and further questions may require entirely new methods or algorithms. Second, there is the related issue of having numerous available (and somewhat redundant) options for extremely complex NGS bioinformatics data analysis tools.

Because NGS data and their formats frequently change, the analytical tools must adapt; new tools arise frequently through efforts to improve upon initially developed algorithms, or to complement other methods. One can often identify dozens or even hundreds of individual tools that can perform the same type of analysis, and it has been an increasing challenge to decide which tools are best for which specific applications. In addition, some tools are tailored to specialized hardware architectures. Lastly, few laboratories have the degree of expertise required to implement robust methods, install the appropriate tools, or construct standardized pipelines for processing data. The need for such expertise can delay studies and make comparisons of disparate studies very difficult. While some systems have allowed the open-source integration of selected tools within a single environment (e.g., Galaxy (Blankenberg et al. 2010)), users must often already know which tools or pipelines to select and what specific parameters to use for their particular goals. A more costly approach includes commercial packages that can perform similar operations and further help to visualize results, but these packages use proprietary software that can be inflexible or, if one does not know the details of the programs and parameters, can affect downstream interpretation.

Because we view bioinformatics as the key bottleneck in the use of NGS data, we present an integrated platform toward Empowering the Development of Genomics Expertise (EDGE). This bioinformatics effort is intended to truly democratize the use of NGS for exploring novel genomes and metagenomes. We developed EDGE Bioinformatics as an initial suite of pre-configured bioinformatics workflows that allow rapid analysis of NGS data, coupled with visualization and interactive features. These features allow users to view results and explore ongoing data processing within an intuitive and user-friendly web-based environment. The software is freely available (https://lanl-bioinformatics.github.io/EDGE/) and a webserver is provided (https://bioedge.lanl.gov/) for use with publicly available data via the NCBI Sequence Read Archive.

## RESULTS

### The EDGE Bioinformatics overview

An overview of the EDGE Bioinformatics workflow is shown in Figure 1, with a more detailed workflow shown in Supplementary Figure S1. Because most sequencers can now output data as one or more FASTQ files we opted for this format (full or compressed) as the required input for raw sequencing data. EDGE can use files derived from multiple libraries, runs or lanes by specifying the location of one or more FASTQ files or by retrieving them from the Sequence Read Archive (SRA) at NCBI (Supplementary Figure S2). EDGE was originally designed for use with Illumina^®^ reads and performs best with these short sequence data types, but the development of alternative workflows are envisioned for future versions to better handle other types of data (e.g. longer reads, different error models, etc.). There are a number of additional options such as specifying number of CPUs to use or allowing batch submission of many samples.

**Figure 1.**
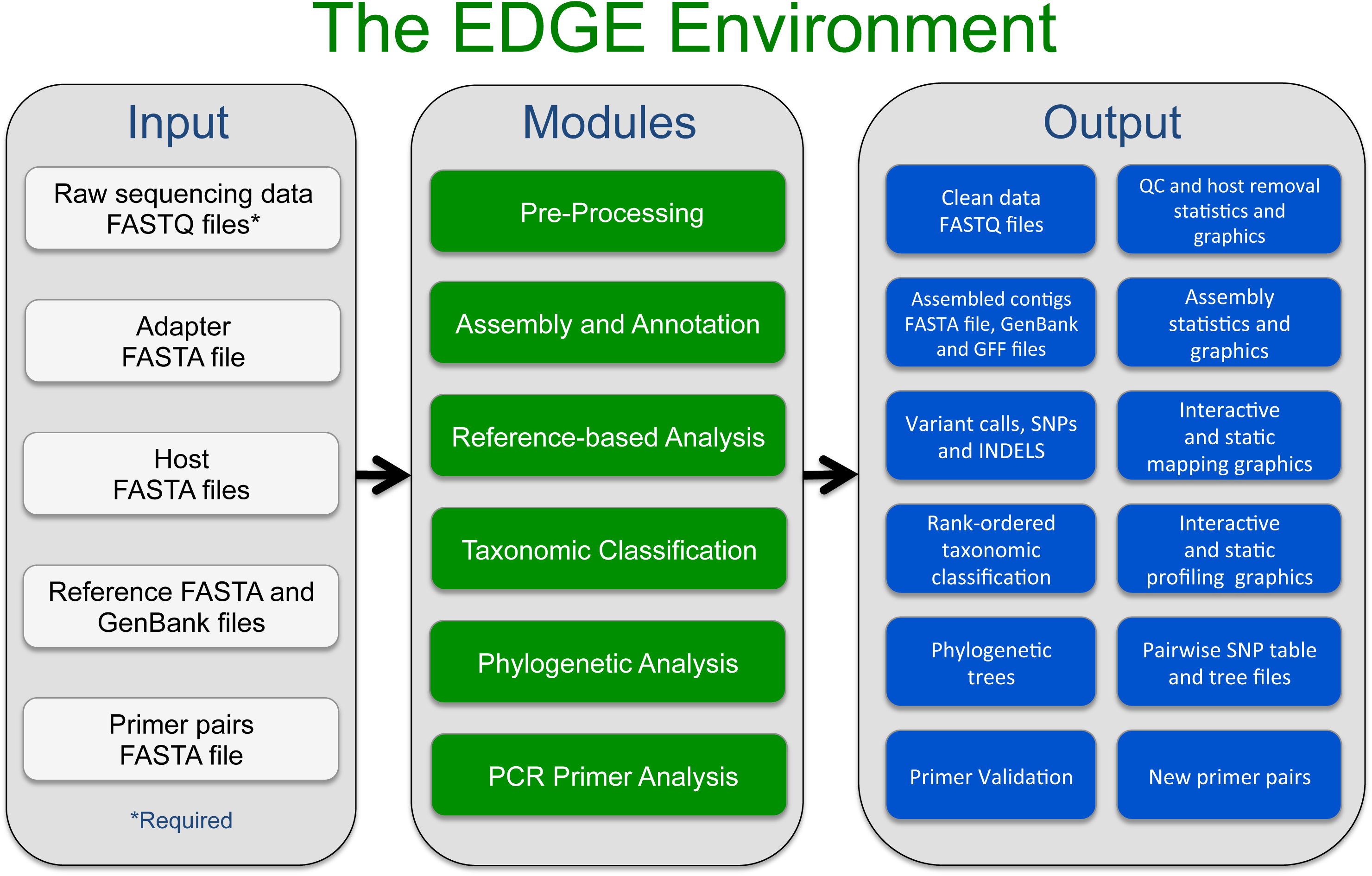
An overview of the EDGE Bioinformatics Environment. The only Inputs required from the user are raw sequencing data and a project name. The user can create specific workflows with any combination of the modules. In addition, tailored parameters dictating how each module functions can be modified by the user. EDGE outputs a variety of files, tables and graphics which can be viewed on screen or downloaded. A more detailed overview is shown in Supplementary Figure S1. All Modules are described in the Methods section.

Optional inputs depend on the selected modules (see Methods) and can include an adapter FASTA file for adapter filtering, a host FASTA file for removal of host reads, PacBio/Nanopore long read FASTA/FASTQ files for use with the SPAdes (Bankevich et al. 2012) assembler, one or more reference genomes for comparative genomic analysis, and a primer pair(s) file in FASTA format for *in silico* primer validation. While there are several optional environmental parameters that can control the way EDGE runs, the users need only specify a project name, select the input file(s), toggle which modules they would like to use, and click Submit. The results of each project are displayed within its own project page (see Methods and Supplementary Figure S3). Descriptions of all modules are in the Methods section.

### Analysis in EDGE

To demonstrate the utility and versatility of EDGE, we tested this platform using a number of different samples that represent varied scenarios, including examples of isolate sequencing and analysis of several clinical metagenome samples with known, suspected, and unknown etiologic agents (Table 1). Not all results are described in depth, but the different datasets are used to highlight some of the various modules and analytic capabilities encompassed within the EDGE Bioinformatics platform. All datasets and project pages with full results are publicly available on our webserver (https://bioedge.lanl.gov/). There, users can view or select and run their own analyses of these data or other publicly accessible SRA data.

**Table 1.** Descriptions of samples and EDGE modules tested.

| Sample description | Sample type (material) | # of reads (millions) | Sequence type | EDGE Modules* |  |  |  |  |  | CPUs | Run Time (hours) |
| --- | --- | --- | --- | --- | --- | --- | --- | --- | --- | --- | --- |
|  |  |  |  | 1 | 2 | 3 | 4 | 5 | 6 |  |  |
| *Bacillus anthracis* strain SK-102<br>SRR1993644 | Isolate (gDNA) | 28.6 | HiSeq 2x101 nt | X | X | X | X | X | X | 8 | 4:12:03 |
| *Bacillus anthracis* strain SK-102<br>SRR1993644 | Isolate (gDNA) | 28.6 | HiSeq 2x101 nt | X | X | X | X | X | X | 20 | 3:33:52 |
| *Yersinia pestis* strain Harbin 35<br>SRR1993645 | Isolate (gDNA) | 15 | GAll 2x110 nt | X | X | X | X | X | X | 8 | 3:35:39 |
| Human Microbiome Project (staggered mock community)<br>SRR172903 | Metagenome (DNA) | 7.93 | GAll 75 nt | X | X |  | X |  |  | 8 | 0:53:59 |
| Patient plasma sample 2014 *Ebola* outbreak (IDBA assembly)<br>SRR1553609** | Metagenome (RNA) | 0.93 | HiSeq 2x100 nt | X | X | X | X |  |  | 12 | 0:38:07 |
| Patient plasma sample 2014 *Ebola* outbreak (SPAdes assembly)<br>SRR1553609** | Metagenome (RNA) | 0.93 | HiSeq 2x100 nt | X | X | X | X |  |  | 12 | 0:47:24 |
| Patient fecal sample 2011 *E. coli* outbreak<br>SRR2164314 | Metagenome (DNA) | 273 | HiSeq 2x100 nt | X | X |  | X |  |  | 8 | 34:43:30 |
| Patient nasal swab acute respiratory illness<br>SRP062772** | Metagenome (DNA) | 2.52 | MiSeq 2x300 nt | X | X |  | X |  |  | 8 | 0:20:59 |

\* EDGE Modules are described in Methods: 1. Pre-Processing; 2. Assembly and Annotation; 3. Reference-Based Analysis; 4. Taxonomic Classification; 5. Phylogenetic Analysis; 6. PCR Primer Analysis
\*\* These samples were retrieved directly from the NCBI SRA.

### Analysis of isolate genome sequencing projects

To highlight and validate some of the features and integration of utilities within EDGE, we tested the various modules using two datasets (sequenced at two different institutions) from recently completed isolate genome sequencing projects: *Bacillus anthracis* strain SK-102 (Johnson et al. 2015b) and *Yersinia pestis* strain Harbin 35 (Johnson et al. 2015a). After quality control, 96-98% of the reads were retained for *B. anthracis* and *Y. pestis* (Supplementary Figure S4). Results from the Assembly and Annotation module were consistent with known genome complexity (repeated elements such as insertion sequences and rRNA operons), genome size, and associated number of genes. The *B. anthracis* assembly was 5.5 Mb in size, consisting of 89 contigs with a maximum contig size of 450kb and an average contig fold coverage of 328X, consistent with the amount of data sequenced (Supplementary Figure S5). The *Y. pestis* assembly (4.6 Mb with 306X fold coverage) was more fragmented (329 contigs) with smaller contig sizes (maximum contig size of 115kb) owing to the large number of repeat sequences within the genome. However, using the reference-based analysis module, all of the *Y. pestis* contigs, and all but a single contig of the *B. anthracis* assembly, could be mapped to the selected reference genome (*Y. pestis* CO92 and *B. anthracis* Ames Ancestor, respectively). More than 98% of the reads of either sample could also be mapped, covering >97-100% of the reference chromosomes and plasmids (Supplementary Figure S6).

While the identities of the organisms sequenced in this case are not in question, the taxonomy classification module can be used to identify a contaminant, or otherwise suggest similarity to another taxon. The consensus for all the taxonomy classification tools encompassed in EDGE confirmed the presumed identities of the organisms sequenced. With *Y. pestis*, both GOTTCHA (Freitas et al. 2015) and Metaphlan (Segata et al. 2012) provided the cleanest results, suggesting only *Y. pestis* reads comprise the dataset (Figure 2A), however with *B. anthracis*, a number of different organisms were found by these tools (Figure 2B), even at the genus level. At the species level, both GOTTCHA and Metaphlan identified *B. cereus* and *Francisella philomiragia* in addition to the dominant *B. anthracis*. In addition, GOTTCHA found signatures of *Y. pestis* and *B. weihenstephanensis*, while Metaphlan suggested *B. thuringiensis* was present. Upon further investigation, we discovered that the *B. anthracis* SK-102 sample was sequenced within the same Illumina lane as many other samples, including *F. philomiragia* ATCC25018, two *Y. pestis* strains (771 and 790), *B. cereus* BACI291, *B. mycoides* BACI084 (a near neighbor to *B. weihenstephanensis* (Soufiane and Cote 2013)), and several fecal samples from Condors (found to contain dominant amounts of *Clostridia* sequences, consistent with dominance of *Clostridia* in the Vulture hindgut (Roggenbuck et al. 2014)). Therefore these additional identifications are likely the result of index cross contamination (or other mis-assignment) of barcodes to sample, often found among samples run within the same lane (Kircher et al. 2012). In addition, and consistent with the bacteria in this sample, GOTTCHA viral analysis suggested three *Bacillus* phages as well as *Staphylococcus* phage SpaA1, which is similar to *Bacillus* prophages and can infect *Bacillus* spp. (Swanson et al. 2012).

**Figure 2.**
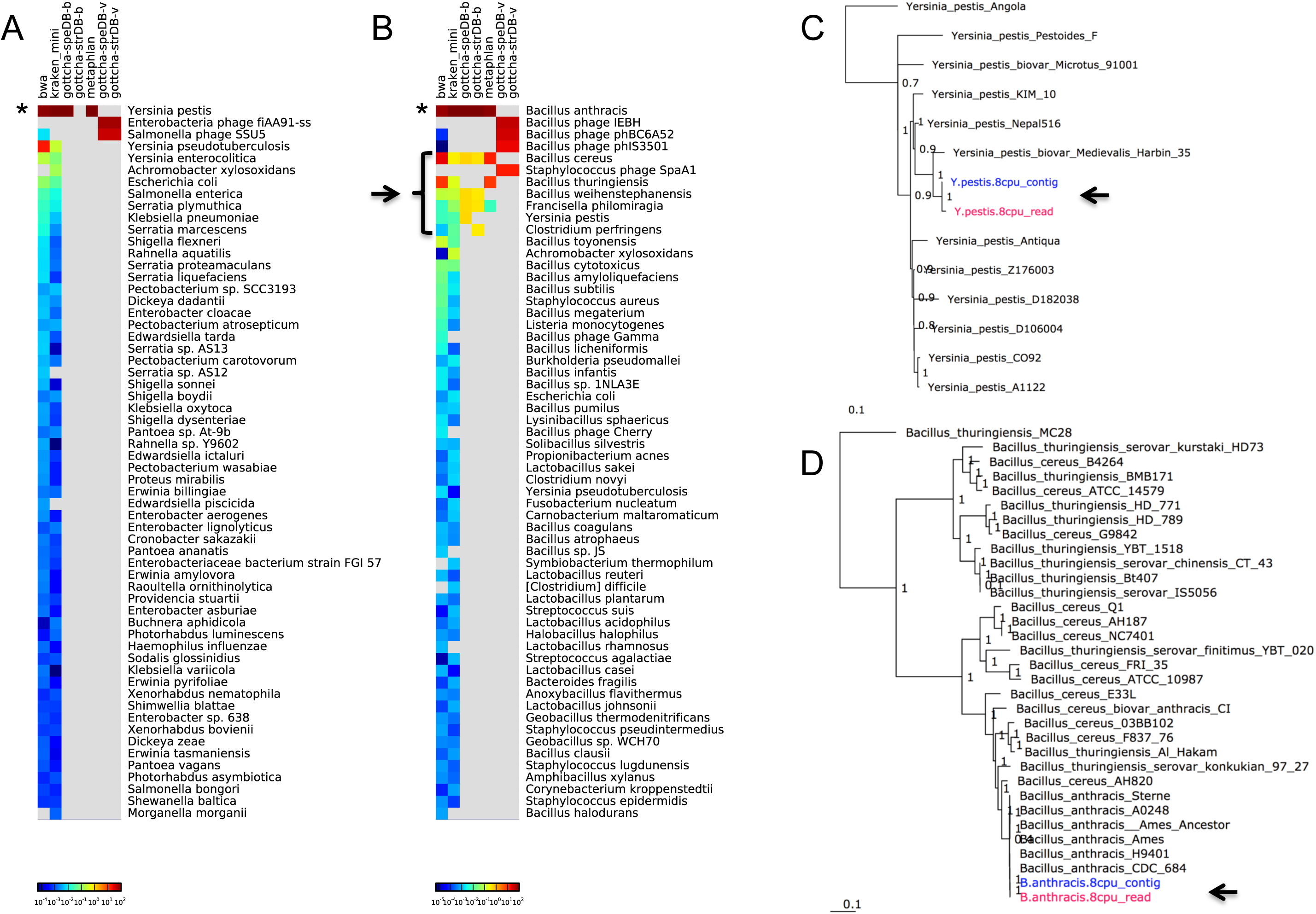
Taxonomy and phylogenetic evaluations of bacterial isolates. Panels A and B show taxonomic classification of reads for A) the *Y. pestis* Harbin35 sample and B) the *B. anthracis* SK-102 sample. The stars indicate the consistent dominant taxonomic calls for all tools, while the black arrow and bracket indicate identified contamination in the *B. anthracis* sample. Panels C and D indicate the inferred phylogenetic trees for the C) *Y. pestis* and D) *B. anthracis;* black arrows point to the read dataset (pink) and contigs (blue) that were placed in these trees.

Phylogenetic analysis was performed for each dataset, selecting all available NCBI RefSeq genomes for either *Y. pestis*, or for *B. anthracis*, *B. cereus*, and *B. thuringiensis*. This phylogenetic module, based on PhaME (Ahmed et al. 2015), independently treats the input reads and resulting contigs (when assembly is selected) for whole genome SNP analysis, and consistently placed the datasets within their respective phylogenetic trees (Figure 2C-D). The *Y. pestis* tree was inferred from a 4.0 Mb core genome with 2,077 SNPs and the *Y. pestis* sample was placed nearest a previously sequenced *Y. pestis* Harbin35. The *Bacillus* tree was based on a core genome of 3.1 Mb with 384,568 SNPs, is fully consistent with known *Bacillus* relationships (Soufiane and Cote 2013), and placed the reads and the resulting contigs of the *B. anthracis* SK-102 closest to *B. anthracis* CDC684.

Using the PCR Primer Tools module, published primers that have been used to detect either *Y. pestis* (Hinnebusch and Schwan 1993; Begier et al. 2006) or *B. anthracis* (Fasanella et al. 2003; Francy et al. 2009; 2012) were input for validation against these isolates and confirmed the appropriate amplicon sizes using electronic PCR against the respective assemblies. For *B. anthracis*, two novel PCR primer pairs were suggested by the primer design software that would specifically amplify only this strain compared with all other NCBI genomes (Supplementary Figure S7).

### Analysis of a mock human microbiome sample of known complexity

The Human Microbiome Project’s (HMP) staggered mock community (Human Microbiome Project 2012) was used to evaluate the metagenome analysis potential of EDGE. This dataset, consisting of sequencing reads derived from a mixture of 21 known bacterial strains and one eukaryotic strain, was analyzed using the Pre-processing, Assembly, and Taxonomy classification modules with default parameters. The FaQCs (Lo and Chain 2014) quality control pipeline retained 81.2% of the reads and 76.7% of the data from the 7.9M read dataset, while the subsequent assembly produced 13,097 contigs totaling 14.8 Mb. Read mapping validation suggested that the assembly represents 77.6% of the reads with a contig average fold coverage of 24X (Supplementary Figure S8). Both the read-(Figure 3A), and contig-based (Figure 3B) taxonomy classification tools accurately identified most of the known community members of this sample with the exception of the eukaryote since these tools were implemented to identify bacteria and viruses. The contig plot of average G+C (%) versus average fold coverage can also help distinguish groups of contigs that belong to different organisms (Figure 3C). Similar graphics and results can be found at various taxonomic levels.

**Figure 3.**
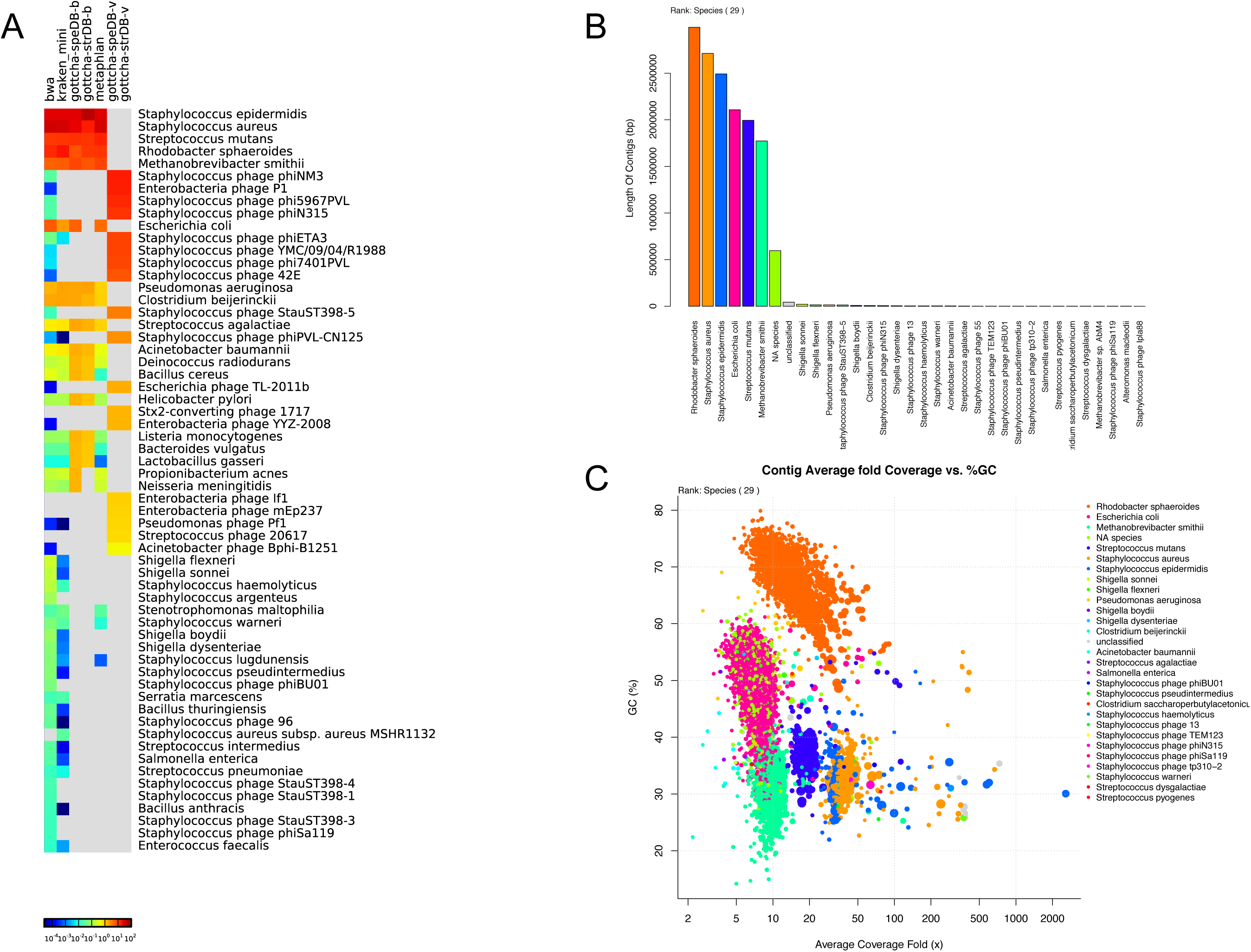
Taxonomic Classification of the HMP staggered mock sample. A) Read-based classification using various taxonomy profiling tools; B) contig-based classification displaying length of all classified contigs per taxon; and C) a scatterplot of contig % GC vs. fold coverage of the contigs, colored by taxon.

### Analysis of complex clinical samples

We also used EDGE to evaluate datasets from several clinical samples with suspected pathogens. In the first example we used EDGE to characterize one of the recent 2014 Ebola outbreak samples. Using the Sierra Leone human plasma RNA sequencing sample SRR1553609 retrieved directly from the SRA, we ran all EDGE modules with the exception of phylogenetic and primer analyses. Pre-processing removed ~25% of the data, and human host removal only identified 605 reads that matched the human reference. IDBA (Peng et al. 2012) assembly of the remaining reads resulted in 1588 contigs, a total assembly size of 665kb and a largest contig of 14.6kb. Due to the complexity of the sample, only 15% of the data assembled. We examined the use of the alternate assembler, SPAdes (Bankevich et al. 2012), with this sample and found an increased run time (Table 1) balanced by an improved 36% read incorporation (vs. 15%) into the assembly, resulting in 12,105 contigs, a total assembly size of >3.8Mb and a largest contig of 18.6kb. Using as reference the *H.sapiens*-wt/GIN/2014/Makona-Gueckedou-633 Zaire ebolavirus (a sequence from Guinea, 2014), we found that only 3,228 reads (0.43% of the input reads) could be mapped to the genome, covering 98.9% of the length with 10 potential single nucleotide variants. Two of the IDBA contigs overlapped and together covered 99.2% of the genome, while a single SPAdes contig covered 97.8% of the reference. Both assemblies identified the same 8 SNPs with respect to the reference genome. The genome browser in EDGE helped resolve the disparate variant analysis found between the reads and the contigs (Figure 4). While almost all of the reads confirmed all 8 SNPs found within the contigs, the two additional variants identified with read-based analysis likely reflected the quasispecies nature of the virus, with strong support but fewer than 50% of the reads at those positions carrying the additional point mutations. This shows the utility of a multi-pronged approach when performing such comparisons. The taxonomy classification module showed that Ebola could indeed be found within the reads, though only with the GOTTCHA and BWA pipelines, and also provided a list of bacteria that may have also been present within the sequenced sample, including *Ralstonia*, *Bradyrhizobium*, *Propionibacterium*, and *Pseudomonas* (Supplementary Figure S9). The contig-based taxonomy analyses also clearly showed Ebola virus to be present, and confirmed that many contigs belonged to the same bacterial groups identified by read-based analyses.

**Figure 4.**
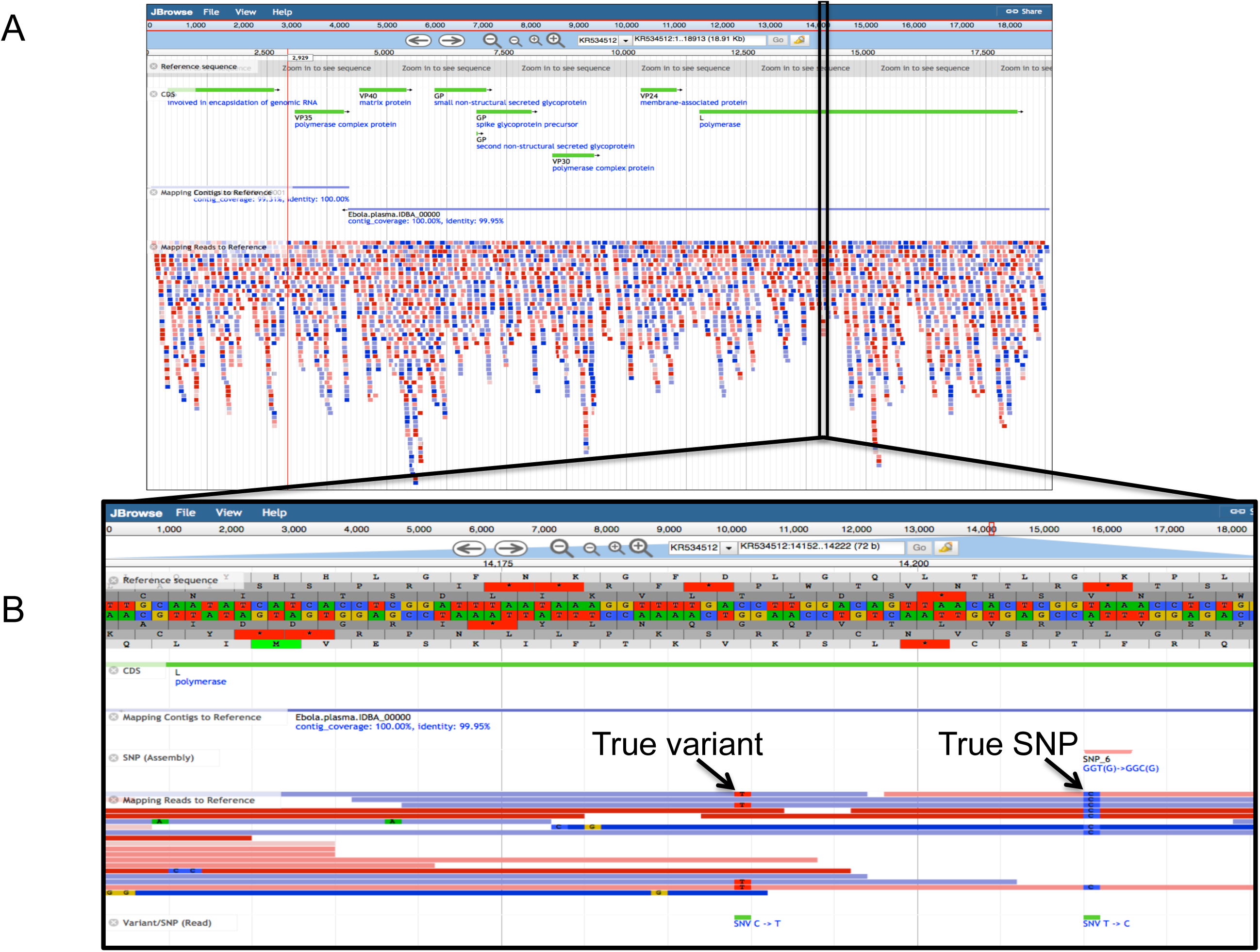
Interactive genome browsing view of a reference-based analysis in EDGE with a human clinical sample containing Ebolavirus. A) An Ebola reference genome and its genes (green lines) are displayed together with contig-based (using IDBA) and read-based comparisons. The two contigs (blue lines) from IDBA are shown aligned along the length of the reference as well as the reads (red and blue). B) A zoomed-in view of one section of the genome where SNPs were identified. The SNP and coding difference is outlined under the contig alignment, while the variants are indicated under the read alignments.

In the second clinical example, we analyzed data derived from a fecal sample of a patient returning from Germany during the 2011 enterohemorrhagic *Escherichia coli* outbreak, and who was suspected of harboring *E. coli* O104:H4. Trimming and filtering removed 13.3% of the bases while host removal identified only 0.15% of the reads as human and 0.02% as PhiX (a spike-in control commonly used in Illumina sequencing). Assembling the remaining 253M reads resulted in 2,957 contigs totaling 10.5 Mb, comprising 23.9% of the reads. The single chromosome and three plasmids of *E. coli* O104:H4 2011C-3493, were used as reference for both read-and contig-based comparisons. Using reads, 99.99% of the reference chromosome was covered at 115X, while the three plasmids were covered 100% at fold-coverages ranging from 250X for the largest plasmid to 7.6 million fold coverage for the smallest plasmid. Using contigs, all replicons were covered >99.7% with the exception of the small plasmid which was absent from the assembly (this absence is likely due to the excessive fold coverage known to create assembly issues). All taxonomy profiling tools clearly showed that *E. coli* (or *Shigella*) was the dominant organism and that the Shiga-toxin phage was also present (Supplementary Figure S10). Whole genome SNPs were identified and phylogenetic analysis was performed with both reads and contigs, easily done within EDGE using the drop down menu to select 68 *E. coli* and *Shigella* genomes. Both the predominantly *E. coli* metagenome reads and the assembled contigs were placed within the same clade as the other *E. coli* O104 strains, reaffirming the initial suspicion of *E. coli* O104:H4 as the etiologic agent (Figure 5A).

A nasal swab sample from a patient with acute respiratory illness of unknown etiology was used as a final test of EDGE’s utility for analysis of clinically derived metagenomic datasets. In this case, while >99% of the data passed FaQCs quality control, the majority of sequence reads (78.9%) were human-derived and removed (data not shown). The remaining reads were submitted to SRA and used for assembly and taxonomy classification. A number of expected organisms (Rawlings et al. 2013; Bassis et al. 2015) ranked among the most abundant genera identified, including *Prevotella, Veillonella* and *Streptococcus*. Unexpectedly, *E. coli* was identified by GOTTCHA, and also detected (at a substantially lower level) by BWA and Kraken mini (Figure 5B). Upon closer inspection, the mapping results demonstrated that all of the *E. coli* hits were to the plasmid (with no matches to the chromosome) in *E. coli* strain ABU83972, covering approximately 80% of this replicon. Interestingly, this plasmid is very similar (>90% identity) to a number of enteric plasmids, as well as to the *Corynebacterium renale* plasmid pCR1, suggesting that the presence of this plasmid might be the result of colonization or infection by a *Cornyebacterium* species, which are common in nasal cavities (Bassis et al. 2015). This hypothesis is partially supported by BWA and Kraken, which identified a different *Cornyebacterium* at low levels, as well as by 16S sequence data in which *E. coli* is not detected but the genus *Cornyebacterium* is found (Supplementary Table S1). As a result of these findings a new feature now present in EDGE separates plasmid from chromosomal hits for GOTTCHA, thereby allowing for greater specificity in evaluating taxonomic profiling results (Figure 5C). The differences in bacterial species found by Metaphlan compared with all other tools can be explained by the additional draft genome references included within the Metaphlan database (Segata et al. 2012) and which are not yet available in RefSeq.

**Figure 5.**
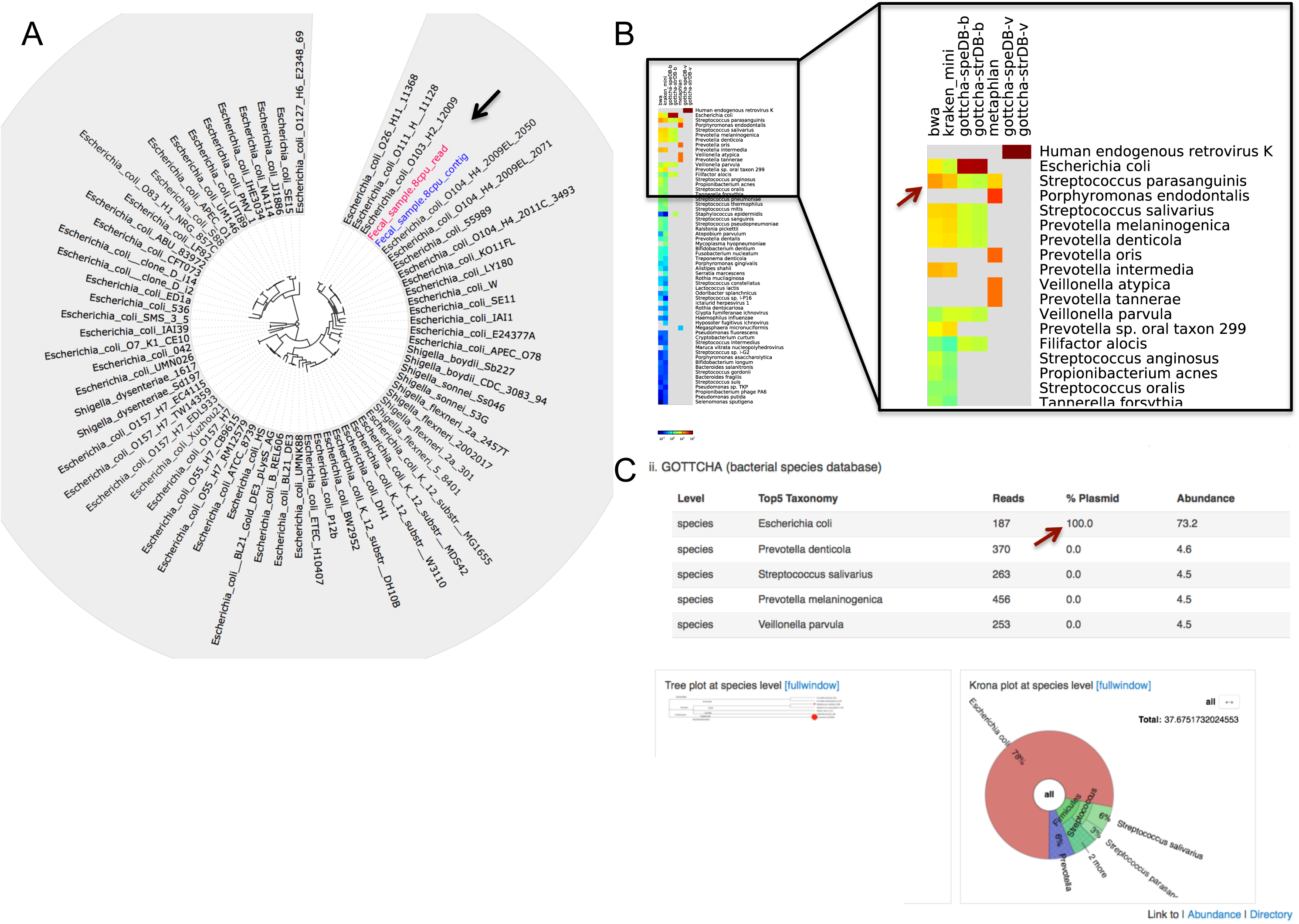
Phylogenetic and taxonomic analysis of human clinical samples with suspected and unknown causative agents. A) Circular phylogenetic tree clearly places within the *E. coli* 0104 group both the raw reads and the contigs obtained from a clinical fecal sample. B) A comparative heatmap view of identified taxa from a nasal swab sample demonstrates the abundance of typical nasal cavity organisms. C) The *E. coli* identified with GOTTCHA in the nasal swab sample (in B) is described in greater detail under the tool-specific EDGE view (red arrow), showing the percent of hits to plasmids for each identified taxon; below are a taxonomic dendrogram featuring the taxa detected with circles representing relative abundance, and a Krona plot view of the same data.

## DISCUSSION

As the number of investigations that apply sequencing continues to climb, the wider genomics community will greatly benefit from a user-friendly bioinformatics environment of integrated tools and pipelines designed to address a large number of scenarios and scientific end-goals. The initial system and the tools we developed and used in EDGE are available as open source software, and we encourage other developers to contribute best-practice tools and pipelines, as there are yet a number of use cases not addressed within this initial platform. For the tools in current use, the focus was on accuracy, speed, flexibility, and ability to run within a modest computational environment. In some cases, like with read-based taxonomy profiling, given that this is a still emerging field of exploration, we provide a suite of tools based on different algorithms, and present a comparative view of the results for further scrutiny by researchers. In other cases, tools were selected that perform well under a diverse set of circumstances, and are computationally friendly with respect to speed and memory considerations. While novel tools continue to be developed and databases continue to grow, future focus will be on the systematic incorporation of better tools and updating of databases alongside the development of new modules.

Collectively, our results and experiences suggest that EDGE provides significant advantages over the current status quo. For example, significant expertise is generally required to determine what tools (and parameters) are ideal for different scenarios, and in many cases, to run these tools and manipulate the results. EDGE assists non-expert users by providing pre-defined pipelines to run cutting-edge tools and a web interface that makes inspection of results quick and easy. Comparative views of results output by complex metagenome taxonomy profiling tools distinguish this system from all others along with the ability to easily perform whole genome SNP phylogenies with user-selected genomes. The ability to integrate read-based with assembly-based analyses provides complimentary views of genomic data. Real time tracking of projects and system resources allows for better monitoring and job queuing. With embedded log files detailing the specifics of each run, a wide adoption of systems like EDGE can also provide a form of standardized data analysis which would allow for more robust comparisons to be made across different independent projects and laboratories.

EDGE is a unique bioinformatic software package both for the variety of open-source tools that are encompassed and for its ease of use. To our knowledge, there is no other freely available bioinformatic software package that incorporates these types of analyses and tools within a framework of intuitive pipelines and interactive graphical and tabular results. This software package is designed to enable scientists with limited experience in bioinformatics to perform a variety of genomic analyses with resources that are available in smaller laboratories, rather than requiring extensive computational and personnel infrastructure. The EDGE Bioinformatics software therefore represents a critical step forward in democratizing genomics analyses.

## METHODS

### EDGE Bioinformatics computational design

EDGE Bioinformatics is built around a collection of publicly available, open-source software packaged in six modules. The main wrapper script is written in Perl, while the various tools currently include BLAST, version 2.2.26 (Altschul et al. 1990), BowTie2, version 2.1.0 (Langmead and Salzberg 2012), BWA, version 0.7.9 (Li and Durbin 2010), FaQCs, version 1.33 (Lo and Chain 2014), FastTree, version 2.1 (Price et al. 2010), GOTTCHA, version 1.0b (Freitas et al. 2015), IDBA_UD, version 1.1.1 (Peng et al. 2012), SPAdes, version 3.5.0 (Bankevich et al. 2012), JBrowse, version 1.11.6 (Skinner et al. 2009), jsPhyloSVG, version 1.55 (Smits and Ouverney 2010), Kraken, version 0.10.4-beta (Wood and Salzberg 2014), KronaTools, version 2.4 (Ondov et al. 2011), MetaPhlAn, version 1.7.7 (Segata et al. 2012), MUMmer3, version 3.23 (Kurtz et al. 2004), Phage_Finder, version 2.1 (Fouts 2006), PhaME (Ahmed et al. 2015), Primer3, version 2.3.5 (Untergasser et al. 2012), Prokka, version 1.11 (Seemann 2014), RATT, version 08-Oct-2010 (Otto et al. 2011), RAxML, version 8.0.26 (Stamatakis 2014), and SAMtools, version 0.1.19 (Li et al. 2009).

All tools and modules can be run on Unix command line, however we provide a user-friendly web-based graphic user interface (GUI). The GUI is primarily implemented using the JQuery Mobile javascript framework and HTML5 on the client-side, and implements perl CGI using Apache or Python on the server-side. This implementation makes EDGE accessible on any platform, including all smartphones, tablets, and desktop devices. The EDGE software tools were selected or developed based on the desire (and need) for both accuracy and speed, with the assumption of moderate computational hardware resources. More detail regarding the installation, implementation, and the tools encompassed within EDGE can be found at http://edge.readthedocs.org/.

The modular design and open source license also allow other researchers to expand the available capabilities beyond our initial implementation. For expert bioinformaticians, another benefit is that EDGE can also be integrated into other workflows and be used via command line to submit jobs on a cluster. More information can be found at the EDGE homepage (https://lanl-bioinformatics.github.io/EDGE/), and the software is available at https://github.com/LANL-Bioinformatics/edge. We also offer a partially modified, public EDGE webserver, available at https://bioedge.lanl.gov/, which can be used to analyze publicly available data deposited in the NCBI SRA or EMBL ENA. All project datasets and results discussed in this manuscript are provided on this site.

One of the key features of the EDGE Bioinformatics platform is that visualization of the results is fully integrated with, and accessible directly on, the webpage in real time. Many graphics are displayed on each project page as thumbnails that link to either a full-page view or a lightbox (quick zoom) view, including quality control graphics, assembly summary charts, heat maps, phylogenetic trees, etc. In addition, there are links to the interactive genome browser JBrowse and to interactive classification results via Krona, as well as links to output directories where all resulting data for each pipeline are stored.

Because some of the most challenging aspects of genomics involve the exponentially increasing size of datasets and the resources required to move large datasets, a key benefit of the EDGE Bioinformatics software is that it can be implemented on a stand-alone server that can access datasets in local storage or in network-mounted space. We have tested EDGE Bioinformatics with datasets of up to hundreds of millions of reads, on a variety of servers (e.g. 12-64 core servers with 64GB-512GB of RAM), with run times ranging from minutes to hours. Using more CPUs will decrease runtime (see Table 1). All projects run for this manuscript were performed on the publicly available server (https://bioedge.lanl.gov/), which is a Dell PowerEdge R720 with 24 cores, 512GB RAM, and 7 TB disk space. On this particular server, we have restricted use to a maximum of 20 CPUs that can be specified by any given analysis.

A user management system has been implemented to provide a level of privacy/security for a user’s submitted projects. When this system is activated, any user can view projects that have been made public, but other projects can only be accessed by logging into the system using a registered local EDGE account or via an existing social media account (Facebook, Google+, Windows, or LinkedIn). The users can then run new jobs and view their own previously run projects or those that have been shared with them.

### The project page layout

A left navigation menu on the EDGE website provides access to the Home page, the Run EDGE page (to initiate a new project) and the Projects list, allowing users to navigate to any desired project page (Supplementary Figure S3). A page for each project is produced as soon as it is launched within EDGE and allows the user to monitor the progress of the run and access the output summaries of each pipeline as they complete in real time. Each project page provides a summary of the project, and under a ‘General’ tab, a description of the input(s) provided, the modules selected for the run along with their run time statistics, and access to log files, the output directory, and a final PDF report.

A link in the upper right corner provides access to a sliding panel that contains a job progress widget, a resource monitoring widget, and an action widget. Once the job is submitted, the job progress widget reports the status for each analysis step in real time. The resource monitoring widget provides a real time view of the computational system running EDGE, and allows the user to anticipate whether there are sufficient resources to simultaneously run additional jobs, or if some projects should be moved to a different storage location. For example, projects will fail to complete one or more of the modules if there is insufficient storage for the outputs. The action widget provides the user some flexibility over the project, including allowing a user to interrupt, rerun, delete, and move his or her submitted jobs. The user can also share the project with other users, publish the project such that any user can access the results, or make the project private again (unpublish). In addition, there is a command line ‘live log’ view, which displays the real time actions and the Unix commands launched by EDGE.

### The EDGE modules and their outputs

All of the six main modules within the EDGE Bioinformatics environment are optional and can be selectively run as individual modules or in any combination, thus affording the user maximum flexibility in customizing each analysis to particular specifications. These consist of: 1) a preprocessing module that performs quality control, trimming, and removal of sequences matching an unwanted target (e.g. host removal); 2) a *de novo* assembly module which assembles the data, validates the assembly, and annotates the resulting contigs; 3) a reference-based analysis module, which allows users to select one or more references to which reads (and contigs) are compared; 4) a taxonomy classification module, which classifies reads (and contigs); 5) a phylogenetics module, which calculates a core genome, determines all SNPs, and infers a phylogenetic tree from a number of input genomes; and 6) a primer and assay module which allows users to validate *in silico* known primers against the *de novo* assembly, or to design new primers that uniquely amplify short sequences within the *de novo* assembly. The latter module does require an assembly for primer analysis.

Each module comprises a Perl wrapper with one or more bioinformatics tools tailored to handle NGS reads and/or contigs, as well as several scripts to parse and post-process the results. The users can also adjust a limited set of parameters or toggle options within each module. EDGE produces a web page for each project with many different summaries of the results for each module, including the statistics of the run (each module and time to completion), summary log files and a PDF summary of all results, along with more detailed results of each individual module. Each module outputs a number of files, which are accessible via a directory link and are summarized with both text and figures along with some interactive graphics all within the context of the website.

#### Pre-Processing (Supplementary Figure S1, module 1)

This module consists of two independent, selectable pipelines. For data quality control, the FaQCs software is used to analyze all reads for quality and to trim or filter out reads using default parameters, unless these are changed by the user (optional). Using an input reference FASTA, EDGE can also filter unwanted reads that align to a selected reference. While this ‘Host Removal’ function was originally envisioned to exclude host reads when inputting clinical samples or those derived from known animals, this component can remove any data that aligns to the input reference, allowing users to selectively remove any other target genome(s). Some built-in references include the most recently updated GRCh38 Human reference and PhiX, which is often used as a control within Illumina runs. This module aims to provide high-quality, clean reads for any subsequent analysis by EDGE. If this module is not selected, the raw data will be used for all downstream process modules.

Statistics and graphical outputs of the data, prior to and after processing, are provided for user interpretation, along with access to the cleaned data files. The major outputs of this module are shown in Supplementary Figure S1 (A, B and C) and example screen shots of output from the EDGE webpage can be found in Supplementary Figure S4.

#### Assembly and Annotation (Supplementary Figure S1, module 2)

EDGE performs *de novo* assembly with the input reads using either IDBA-UD or SPAdes. Because each of these assemblers performs and combines multiple assemblies, both tools are capable of providing reasonable assemblies from a wide variety of sample types, including isolate genomes, single cell projects, and metagenomes. IDBA-UD is used by default (due to time and memory considerations), and the assembly parameter option for kmer sizes begins with k=31 with a step size of 20, until a maximum kmer size is reached (dependent on the read lengths). When this module is selected, assembly validation is performed by mapping the short read input data to the assembled contigs using Bowtie2. Additionally, the user can select to have the assembly annotated (default behavior) using a modified Prokka tool (for the rapid annotation of prokaryotic genomes), and prophages within microbial genomes are detected using Phage_Finder. If there is an available reference that is sufficiently similar to the target genome assembly, EDGE can also use a modified version of the Rapid Annotation Transfer Tool (RATT) to transfer the annotation from the reference GenBank file (a required input for this step) to the assembly. When SPAdes is selected as the assembler, there exists an additional option to input long read data (PacBio or Nanopore) which can help in gap closure and repeat resolution.

The results of this module include the assembled contigs FASTA file, assembly and assembly validation statistics and graphics, the annotation files (gbk and gff), and an interactive JBrowse implementation, which provides visualization of the contigs and their annotation. The major outputs of this module are displayed in Supplementary Figure S1 (D, E, F and G) and an example screenshot can be found in Supplementary Figure S5.

#### Reference-based Analysis (Supplementary Figure S1, module 3)

When this module is selected, the user must choose one or more reference genomes (FASTA or Genbank formats) to which the reads (and contigs, if assembly was performed) are compared. Reads are aligned to the input reference using BowTie2 and variants are identified using SAMtools. Any regions left uncovered by reads are also identified and reported in text files. Similarly, contigs are aligned to the same reference(s) using MUMmer and the results parsed using Perl scripts to catalogue SNPs and small insertions or deletions (indels), as well as regions within the contigs that may be novel and do not align to the reference. If Genbank reference files are provided, the variants, SNPs, and uncovered regions of the reference are further analyzed to output any affected genes and reports are generated to display whether the changes also contribute to synonymous or non-synonymous substitutions within coding regions. Reads and contigs that do not map to the reference are parsed into separate FASTA/Q files and an option is available to align these reads and contigs to RefSeq for taxonomic identification.

In addition to the output text files, several graphics along with statistics are provided that outline linear coverage of the reference, depth of coverage along the reference, number of variants, as well as percentages of input reads and contigs mapped to the reference. Interactive JBrowse views allow for the display of the reference and associated annotation (genes, rRNAs, etc.), along with detailed views of the aligned reads and contigs, as well as any SNPs or small indels that have been discovered. The major outputs of this module are displayed in Supplementary Figure S1 (G, H, and I), while example outputs can be found in Figure 4 and Supplementary Figure S6.

#### Taxonomy Classification (Supplementary Figure S1, module 4)

Envisioned primarily for use with metagenomic datasets or with novel genomes, this module allows both read-based and contig-based classification (the latter performed if assembly was also selected). For taxonomic classification of the reads, the user can select one or more of several available metagenome tools (currently GOTTCHA, Kraken, and MetaPhlAn) along with BWA, a read mapper used against RefSeq. The default is to run all tools to take advantage of their different strengths, and to provide users with additional information to help interpret their data. Each of these classifiers has its own algorithm and database, parameters for the search, and required input format, all of which are automatically managed within the EDGE platform. The specific output formats of each tool are unified into a common framework to generate the reports/graphs displayed by EDGE. There is also an option to classify only unassembled reads, if assembly is selected and the user desires to only classify unassembled data.

The results of each read-based taxonomy profiling method are summarized in comparative views (heatmap plots and radar charts summarize the top hits of each tool) at the user-selected level of taxonomy (genus, species, strain). Results are also presented in more detail in individual tool-based views with taxonomy tree dendrograms and Krona charts (e.g. Figure 5C) while more detailed outputs can be found within the directory links.

For contig classification, EDGE aligns contigs to NCBI’s RefSeq database using BWA-mem. While contigs can match multiple taxa, each segment within a contig is assigned to a unique taxon based on best hit score. While the total length within all contigs is calculated per taxon, each contig is also assigned to a unique taxon based on linear coverage. Both the total length per taxon (Length barplot) and the number of contigs (Count barplot) assigned to a taxon are reported, along with a scatterplot showing the identity of the contig, its fold coverage by reads, and its G+C content (Supplementary Figure S6). These results are reported at all levels of taxonomy using the last common ancestor algorithm.

The major outputs of this module are displayed in Supplementary Figure S1 (J and K), while example outputs can be found in Figures 2, 3, 5 and Supplementary Figures S9 and S10.

#### Phylogenetic Analysis (Supplementary Figure S1, module 5)

Because phylogenetic analysis is a highly desired feature for many genomic investigations, we utilize a portion of a newly developed tool, PhaME, which provides the ability to infer a whole genome SNP-based tree from completed genomes, genome assemblies, and even from reads. Briefly, contigs and selected genomes are compared with one another to identify conserved segments while ignoring repeated regions, and reads are mapped to one of these genomes to continue the identification of a conserved core genome. The core genome alignment is used to identify all SNPs and FastTree (default, for speed considerations) or RAxML can be used to generate a phylogenetic tree. This module was envisioned for use primarily with isolate genome projects (however metagenomes have also been successfully used), where a target genome comprises the majority of the sequencing data and the user desires to accurately place this target genome within the context of near neighbor genomes. The user is required to select a minimum of two reference genomes to which the reads and contigs (if available) will be added to infer a phylogeny.

The Newick format tree files, core genome FASTA, and SNP statistics are available in the directory link and the phylogenetic trees, generated using jsPhyloSVG, are provided for easy viewing in either rectangular or circular tree formats (Outputs L and M in Supplementary Figure S1). The input sample (reads and/or contigs) is highlighted within the trees. Output examples can be found in Figures 2 and 5.

#### PCR Primer Analysis (Supplementary Figure S1, module 6)

EDGE also supports both the design and validation of PCR primers based on the assembly. In the validation pipeline, known primers within a user-specified input file are mapped to the assembly using BWA, given a user-defined number of mismatches (default of 1) to determine if an amplicon would be generated. The user can also select a pipeline to design new primers based on the assembly, that will differentiate the input sequenced sample from all other bacteria and viruses in NCBI’s RefSeq database. In this design component, unique regions are identified using BWA, and Primer3 is used to select primer pairs. All primers are further filtered by melting temperature (T_m_) difference to the nearest neighbor background, within a user-specified value (5°C by default).

For primer validation, the primer binding location(s) and product sizes are reported for any submitted primers (output N in Supplementary Figure S1). For primer design, a full list of primers that uniquely amplify a product within the assembled contigs is reported (only five are displayed by default on the project page), along with information on the nearest neighbor amplicon (output O in Supplementary Figure S1). Examples of output for both primer validation and primer design can be found in Supplementary Figure S7.

## DATA ACCESS

All data have been deposited to NCBI and accession numbers are shown below.

*Bacillus anthracis* strain SK-102, **SRR1993644**

***Yersinia pestis* strain Harbin 35, SRR1993645**

**Patient fecal sample, 2011 *E. coli* outbreak, SRR2164314**

**Patient nasal swab, acute respiratory illness, SRP062772**

## ACKNOWLEDGEMENTS AND DISCLAIMERS

This work was supported by Defense Threat Reduction Agency project CB4026 to NMRC and CB10152 to LANL. We thank our beta test users for their valuable feedback. Many thanks to the LANL Genome Programs group, and in particular to the informatics support team at LANL for their great help and feedback with both sequencing and bioinformatics. We thank Jason Gans for his careful reading of the manuscript and his helpful suggestions. We also thank Gerald Quinnan, Pengfei Zhang, Regina Cer, Cassie Redden, Kenneth Frey, Eugene Millar, and the IDCRP who were involved in production of nasal swab sequence data (metagenome and 16S). The views expressed in this manuscript are those of the authors and do not necessarily reflect the official policy or position of the Department of the Navy, the Department of Defense, the National Institutes of Health, the Department of Health and Human Services, nor the U.S. Government. VPM is a military service member of the U.S. Government. This work was prepared as part of his official duties. Title 17 U.S.C. §105 provides that ‘Copyright protection under this title is not available for any work of the United States Government.’ Title 17 U.S.C. §101 defines a U.S. Government work as a work prepared by a military service member or employee of the U.S. Government as part of that person’s official duties.

